# Genome-scale metabolic network reconstruction of the chloroform-respiring *Dehalobacter restrictus* strain CF

**DOI:** 10.1101/375063

**Authors:** Kevin Correia, Hanchen Ho, Radhakrishnan Mahadevan

**Author notes:** Corresponding author: Radhakrishnan Mahadevan.

## Abstract

**Background:** Organohalide-respiring bacteria (OHRB) play an important role in the global halogen cycle and bioremediation of industrial sites contaminated with chlorinated organics. One notable OHRB is *Dehalobacter restrictus* strain CF, which is capable of respiring chloroform to dichloromethane. Improved bioremediation strategies could be employed with a greater understanding of *D. restrictus*’ metabolism in isolate and community cultures. To this end, we reconstructed the genome-scale metabolic network of *D. restrictus* to study its metabolism in future studies using flux balance analysis.

**Method:** The RAST annotation server and Model SEED framework were used to obtain a draft metabolic network reconstruction. Additional curation was required for its acetyl-CoA sources, the Wood-Ljungdahl pathway, TCA cycle, electron transport chain, hydrogenase complexes, and formate dehydrogenase complexes.

**Results:** *i*HH623 is the first curated genome-scale metabolic model in the Peptococcaceae family. It spans 1087 reactions and 983 metabolites, covering 623 genes (21% of all ORF’s). Its potential sources of acetyl-CoA are pyruvate ferredoxin oxidoreductase, pyruvate formate lyase, acetyl-CoA synthetase, phosphate acetyltransferase, and CO-methylating acetyl-CoA synthase. NADPH may be regenerated by isocitrate dehydrogenase, malic enzyme, NADP-reducing hydrogenase, cytosolic formate dehydrogenase, ferredoxin-dependent bifurcating transhydrogenase, 5-methyltetrahydrofolate dehydrogenase, and 5-10-methylenetetrahydrofolate. Additional reactions that were added or removed to the *D. restrictus* reconstruction are discussed.

**Conclusions:** We reconstructed the genome-scale metabolic network of *D. restricus* by obtaining an initial draft with the RAST server and Model SEED framework. Curation was required for *D. restricus*’ acetyl-CoA sources, TCA cycle, electron transport chain, hydrogenase complexes, and formate dehydrogenase complexes. This metabolic model can be used to decipher *D. restrictus*’ metabolism in isolate and community cultures in future studies, or as a template to reconstruct the metabolic network of other Peptococcaceae species. The extensive curation of the draft metabolic network reconstruction highlights the need to be cautious of automated metabolic network reconstruction.

## 1 INTRODUCTION

Organohalide-respiring bacteria (OHRB) play an important role in the global halogen cycle and bioremediation of industrial sites contaminated with chlorinated organics, such as chloroform, 1,1,1-trichloroethane and 1,1-dichloroethane (Jugder et al., 2016). One notable OHRB is *Dehalobacter restrictus* strain CF, which is capable of dechlorinating chloroform to dichloromethane (Grostern et al., 2010). Improved bioremediation strategies could be employed with a greater understanding of *D. restrictus*’ metabolism in isolate and community cultures. We reconstructed the genome-scale metabolic network of *D. restrictus* CF, with the aim to better understand its metabolism in future studies using flux balance analysis (Orth et al., 2010).

## 2 METHODS

### Draft and curated reconstruction

A schematic for the reconstruction process is outlined in Figure 1. A genome annotation for *D. restrictus* strain CF (accession no. NC 018866) (Tang et al., 2012) was obtained via the RAST server (Aziz et al., 2008), which was used to reconstruct a draft genome-scale metabolic model with the Model SEED framework (Devoid et al., 2013). The reconstructed metabolic network of *D. restrictus* was subsequently validated against its GenBank annotation and an in-house collection of manually-curated sequences. Reactions derived solely from hypothetical proteins, protein domains, or enzymes with ambiguous substrates were excluded. Redundant/lumped reactions were also removed. Genes with metabolic annotations not included in the initial draft reconstruction were reviewed for inclusion using PaperBLAST (Price and Arkin, 2017).

**Figure 1.**
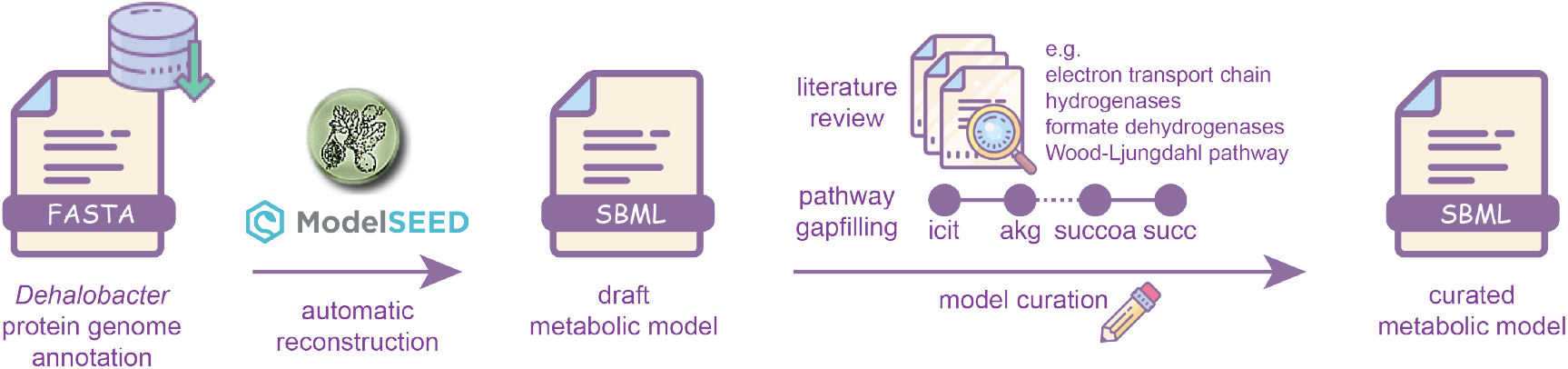
Genome-scale metabolic network reconstruction steps for *Dehalobacter restrictus* strain CF. A genome annotation was obtained from the RAST server, and a draft metabolic network with the Model SEED framework. Curation was required for central metabolism, including the electron transport chain, hydrogenases, formate dehydrogenases, and the Wood-Ljundahl pathway. Pathway gapfilling was also performed to add reactions that were not captured in the draft reconstruction.

### Biomass equation

Reactions to synthesize 1 gram of protein, RNA, DNA, peptidoglycan, and phospholipids from their precursors were added to the reconstruction, rather than having a lumped reaction of precursors to biomass, to allow the biomass composition to be easily manipulated. The amino acid composition of protein from *Bacillus subtilis* was used as the basis for *D. restrictus* (Dauner et al., 2001). The biomass composition was set to 50% protein, 10% RNA, 5% DNA, 5% phospholipid, 25% peptidoglycan, and 5% ash, which is consistent with slow-growing microbes. The growth associated maintenance (GAM) and non-growth associated maintenance (NGAM) were set to 60 mmol · gDCW^−1^ and 19.2 mmol · gDCW^−1^ ·day^−1^, respectively.

## 3 RESULTS & DISCUSSION

The *D. restricus* metabolic model includes 1087 reactions and 983 metabolites, spanning 623 genes (21% of its ORF’s). Model statistics are outlined in Table 1. Compartments include the cytoplasm, periplasm, inner membrane, and the extracellular.

**Table 1.** Model statistics for the curated genome-scale metabolic network reconstruction of *Dehalobacter restrictus* strain CF. The reconstruction captures 21% of its genes.

| Subsystem | Genes | Reactions | Metabolites |
| --- | --- | --- | --- |
| All reactions | 623 | 1087 | 983 |
| DNA polymerase | 16 | 1 |  |
| RNA polymerase | 4 | 1 |  |
| Protein synthesis | 57 | 2 |  |
| Metabolism | 489 | 758 |  |
| Transporters | 43 | 125 |  |
| Exchange reactions | 0 | 69 |  |
| Demand reactions | 0 | 95 |  |

### Acetyl-CoA metabolism

In total there are five sources of acetyl-CoA in *D. restricus*’ metabolic network: pyruvate ferredoxin oxidoreductase, pyruvate formate lyase, acetyl-CoA synthetase, phosphate acetyltransferase, and CO-methylating acetyl-CoA synthase. These reactions expand the solution space and complicate FBA/FVA simulations without additional assumptions. Acetyphosphate acetyltransferase, CO dehydrogenase, and CO-methylating acetyl-CoA synthase from the Wood-Ljungdahl pathway were not part of the draft reconstruction.

### TCA cycle

The TCA cycle in *D. restrictus* is not complete. Succinate dehydrogenase and fumarate reductase genes are absent in *D. restrictus*’ genome (Wang et al., 2017); fumarate reductase was in the initial RAST annotation, but removed in the final reconstruction. Curiously, malate dehydrogenase is also absent in its genome (Wang et al., 2017), preventing the typical bifurcating TCA cycle present in many anaerobes (Amador-Noguez et al., 2010). Ferredoxin-dependent 2-oxoglutarate dehydrogenase from the TCA cycle was added to the reconstruction as it was absent in the RAST annotation.

### Redox metabolism

Possible NADPH sources include malic enzyme, isocitrate dehydrogenase, bifurcating transhydrogenase (NfnAB) with additional promiscuous activities, NADP-reducing hydrogenase (HynABCD), 5,10-methylenetetrahydrofolate dehydrogenase, 5-methyltetrahydrofolate dehydrogenase, and formate dehydrogenase. The electron transport chain, hydrogenases, and formate dehydrogenases reactions all required curation since they were not included in the initial Model SEED reconstruction. Their genes are outlined in Table 2.

**Table 2.** Curated hydrogenase, bifurcating transhydrogense, Complex I, and formate dehydrogenase complexes in the *Dehalobacter restrictus* strain CF metabolic network reconstruction. Localization of the complexes are listed.

| Enzyme | Reaction | Localization | Locus tags | References |
| --- | --- | --- | --- | --- |
| HymABC<br>[Fe-Fe] hydrogenase | 2 Oxidizedferredoxin[c] + H<sub>2</sub>[c] ↔ 2 H<sup>+</sup>[c] + 2 Reducedferredoxin[c] | cytoplasmic | DCF50.p1647-49<br>DCF50.p933-35 | Kruse et al. (2015) |
| EchABCDEF<br>Energy-conserving hydrogenase | 2 Oxidizedferredoxin[c] + H<sub>2</sub>[c] + 2 H<sup>+</sup>[p] ↔ 4 H<sup>+</sup>[c] + 2 Reducedferredoxin[c] | membrane bound | DCF50.p837-42<br>DCF50.p2232-37 | Welte et al. (2010) |
| HupABC(D)<br>[Ni-Fe]-uptake hydrogenase | H<sub>2</sub>[p] + cytochrome-bd-ox[i] → 2 H<sup>+</sup>[p] + cytochrome-bd-red[i] | membrane bound, periplasmic | DCF50.p1849-52<br>DCF50.p1688-90<br>DCF50.p2131-33 | Volbeda et al. (2013) |
| HndABCD<br>NAD(P)-reducing hydrogenase | NAD(P)[c] + H<sub>2</sub>[c] → NAD(P)H[c] + H<sup>+</sup>[c] | cytoplasmic | DCF50.p1597-600 | de Luca et al. (1998) |
| NfnAB<br>Electron-bifurcating transhydrogenase | H<sup>+</sup>[c] + 2 NADP[c] + NADH[c] + 2 Reducedferredoxin[c] ↔ NAD[c] + 2 NADPH[c] + 2 Oxidizedferredoxin[c] | cytoplasmic | DCF50.p511-12 | Huang et al. (2012) |
| NDH1<br>Complex I | NADH[c] + 5 H<sup>+</sup>[c] + Menaquinone 8[i] → NAD[c] + 4 H<sup>+</sup>[p] + Menaquinol 8[i] | membrane-bound, cytoplasmic | DCF50.p2293-303 | Brandt (2006) |
| FdhAEFH<br>Formate dehydrogenase | Formate[c] + NAD(P)[c] ↔ NAD(P)H[c] + CO<sub>2</sub>[c] | cytoplasmic | DCF50.p1622-27 | Kruse et al. (2015) |
| HycABCDEFG<br>Formate hydrogenlyase | Formate[c] + H<sup>+</sup>[c] → CO<sub>2</sub>[c] + H<sub>2</sub>[c] | membrane bound, cytoplasmic | DCF50.p760-66 | McDowall et al. (2014) |
| FdoGHI<br>Formate dehydrogenase-N | Formate[p] + H<sup>+</sup>[c] + Menaquinone 8[i] → CO<sub>2</sub>[p] + Menaquinol 8[i] | membrane bound, periplasmic | DCF50.p923-27 | Wang and Gunsalus (2003) |

### Hydrogenases

There are four types of hydrogenases encoded in *D. restrictus*’ genome: cytoplasmic ferredoxin-dependent [Fe-Fe]-hydrogenase (Hym-type), energy-conserving hydrogenase (Ech-type), [Ni-Fe]-uptake hydrogenase (Hup-type), and NADP-reducing hydrogenase (Hnd-type). Hnd-type was the only hydrogenase present in the draft reconstruction. The electron transfer between Hup-type hydrogenase to reductive dehalogenase has not been fully elucidated (Fincker and Spormann, 2017). For the sake of simplicity, reduced cytochrome b transfers electrons directly to menaquinone in our reconstruction, without any proton motive force. Similar to acetyl-CoA, the presence of multiple hydrogenases complicates flux balance analysis without additional assumptions.

Electron bifurcating transhydrogenase (NfnAB-type) is present in *D. restrictus*’ genome but its activity and regulation are unknown. A characterized homolog has been shown to have various activities, but its dominant activity in *Moorella thermoacetica* is the reversible reduction of NADP via ferredoxin and NADH (Huang et al., 2012). NAD and NADP-dependent ferredoxin reductase were also included in the reconstruction.

### Electron transport chain

Complex I, Hup-type hydrogenase, and energy-conserving formate dehydrogenase are the entry points for electrons into the *D. restrictus* electron transport chain. The initial RAST annotation had ubiquinone as an electron acceptor but was replaced with menaquinone in the final reconstruction. Non-energy-conserving NADH dehydrogenase was removed from the draft reconstruction as there is no strong genomic evidence in *D. restrictus*. All membrane-bound (S)-dihydroorotate dehydrogenase reactions were also removed from the reconstruction; *D. restricus* strain CF’s genome only encodes a cytoplasmic NAD-dependent (S)-dihydroorotate dehydrogenase. The only known terminal electron acceptors in *D. restrictus* are chlorinated organics, via multiple encoded reductive dehalogenase operons (Tang and Edwards, 2013).

### Formate dehydrogenase

*D. restrictus* has two additional formate dehydrogenase reactions: cytoplasmic NAD(P)-dependent formate dehydrogenase, and the membrane-bound formate hydrogenlyase; these enzymes require molybdopterin, which cannot be synthesized by *D. restrictus* (Wang et al., 2017).

All tRNA ligase reactions were added to the reconstruction. Transport and exchange reactions for CO_2_, CO, H_2_, ammonium, chloroform, chlorinated ethenes, malate, pyruvate, glycerol, citrate, oxaloacetate, and all 20 amino acids were included. Demand reactions for all biomass precursors were added to help debug the model.

## 4 CONCLUSION

In summary, we reconstructed the metabolic network of *D. restrictus* to better understand its dechlorinating metabolism in isolate and community cultures. The model includes 623 genes, 1087 reactions, and 983 metabolites. Curation was required for *D. restricus*’ acetyl-CoA sources, TCA cycle, electron transport chain, hydrogenase complexes, and formate dehydrogenase complexes. The presence of multiple reactions involved in the production and consumption of acetyl-CoA, H_2_, and NADPH expand the solution space in flux balance analysis and therefore require additional assumptions to be made with the aide of omics data. The extensive curation in the reconstruction process highlights the need to be cautious of automated metabolic network reconstruction, and the need for improved genome annotation.

## 5 ACKNOWLEDGMENTS

The authors acknowledge Po-Hsiang Wang for his advice on *Dehalobacter restricus*’ metabolism.

This project was supported by the Ontario Genomics SPARK grant.

